# Spatially periodic activation patterns of retrosplenial cortex encode route sub-spaces and distance travelled

**DOI:** 10.1101/100537

**Authors:** Andrew S. Alexander, Douglas A. Nitz

**Affiliations:** University of California, San Diego, Department of Cognitive Science

## Abstract

Traversal of a complicated route is often facilitated by considering it as a set of related sub-spaces. Such compartmentalization processes could occur within retrosplenial cortex, a structure whose neurons simultaneously encode position within routes and other spatial coordinate systems. Here, retrosplenial cortex neurons were recorded as rats traversed a track having recurrent structure at multiple scales. Consistent with a major role in compartmentalization of complex routes, individual RSC neurons exhibited periodic activation patterns that repeated across route segments having the same shape. Concurrently, a larger population of RSC neurons exhibited single-cycle periodicity over the full route, effectively defining a framework for encoding of sub-route positions relative to the whole. The same population simultaneously provides a novel metric for distance from each route position to all others. Together, the findings implicate retrosplenial cortex in the extraction of path sub-spaces, the encoding of their spatial relationships to each other, and path integration.

## Introduction

For most animals, including humans, movement through an environment is frequently constrained to routes along fixed, interconnected pathways. Fluid, efficient navigation under such circumstances demands knowledge of specific routes as well as their locations and orientations within the broader environment. Operationally, a route can often be defined as a series of turns separated by straight-run sections of varying length. As a consequence of the spatial structure associated with such action sequences, a route can also be defined and recognized as a unique space in its own right (i.e. a shape). Notably, identically or similarly shaped routes may exist in many places within the broader environment, potentially at different scales.

The forms of neurophysiological representation associated with route knowledge are only beginning to be understood. Different routes through single environmental positions often yield modulation of in-field firing rates for hippocampal neurons mapping current environmental location^1,2^. Current position within a route itself is represented in the posterior parietal cortex (PPC) and retrosplenial cortex (RSC)^3–6^. However, PPC and RSC route representations diverge. PPC maps position within a route in a manner largely independent of that route’s position in the broader environment, while RSC route encoding is highly sensitive to route position relative to distal cues defining environment boundaries (often referred to as the ‘allocentric’ frame of reference). While neural activity in these regions is also predictive of specific left- and right-turning actions^7–10^, the encoding of progression within a route lies at the scale of the full trajectory since clearly discriminant patterns of activity are generated for different route positions sharing the same action^3,5,11^.

Routes of extensive length and complexity are often described according to their recognizable sub-components. For example, the first leg of a route may be L-shaped followed by a second leg that has an ‘S’ shape. This suggests that there may be regions of the brain, even in rodents, that are capable of representing route sub-components and their relationships to each other. An examination of this possibility at the level of individual neuron spiking dynamics might reveal the form by which encoding of such information is achieved and the rules that govern it.

Among the many structures known to exhibit spatially-specific firing, RSC may be primed to generate the functional dynamics necessary to identify subspaces within a complex route and encode their locations within the environment. RSC is situated as an anatomical intermediary between complementary PPC and hippocampal (HPC) representations of position in a route and position in the environment, respectively. Further, recent work has demonstrated that RSC neuronal ensembles simultaneously encode an animal’s position in multiple spatial frames of reference, including route and allocentric position^5^.

To address the role RSC in sub-route encoding, we trained rats to traverse a route having recursive properties at multiple spatial scales and recorded single-units in RSC. RSC populations generated unique patterns for all route positions. A subset exhibited periodic activation patterns across the full route that oscillated at the scale of specific sub-routes, primarily when they shared the same shape. Further, some RSC neurons oscillated a single time across the full route space, thereby encoding a metric of the animal’s distance from specific locations within the route. Thus, we identify RSC as a region with neurophysiological dynamics that support both the identification and interrelation of sub-spaces within a complex route and provide evidence that RSC can robustly encode distances between all route locations.

## Results

### RSC ensembles generate a unique encoding of all route positions

315 neurons from 5 adult male rats were recorded across all RSC sub-regions during clockwise traversals of a plus-shaped track (**Figure 1A, 1B left;** for histology and waveform discrimination quality see **Supplemental figure 1A-E**). Of these, 229 exhibited a peak in firing rate greater than 3Hz for at least one track position and were not head-direction neurons (**Supplemental figure S1F**). Uninterrupted traversals (**Figure 1B, right**) were identified from tracking data and positional firing rate vectors were generated. Consistent with prior work^5^, maximal firing rates for the full population of neurons were distributed evenly across the full route space (**Figure 1C, left**).

**Figure 1.**
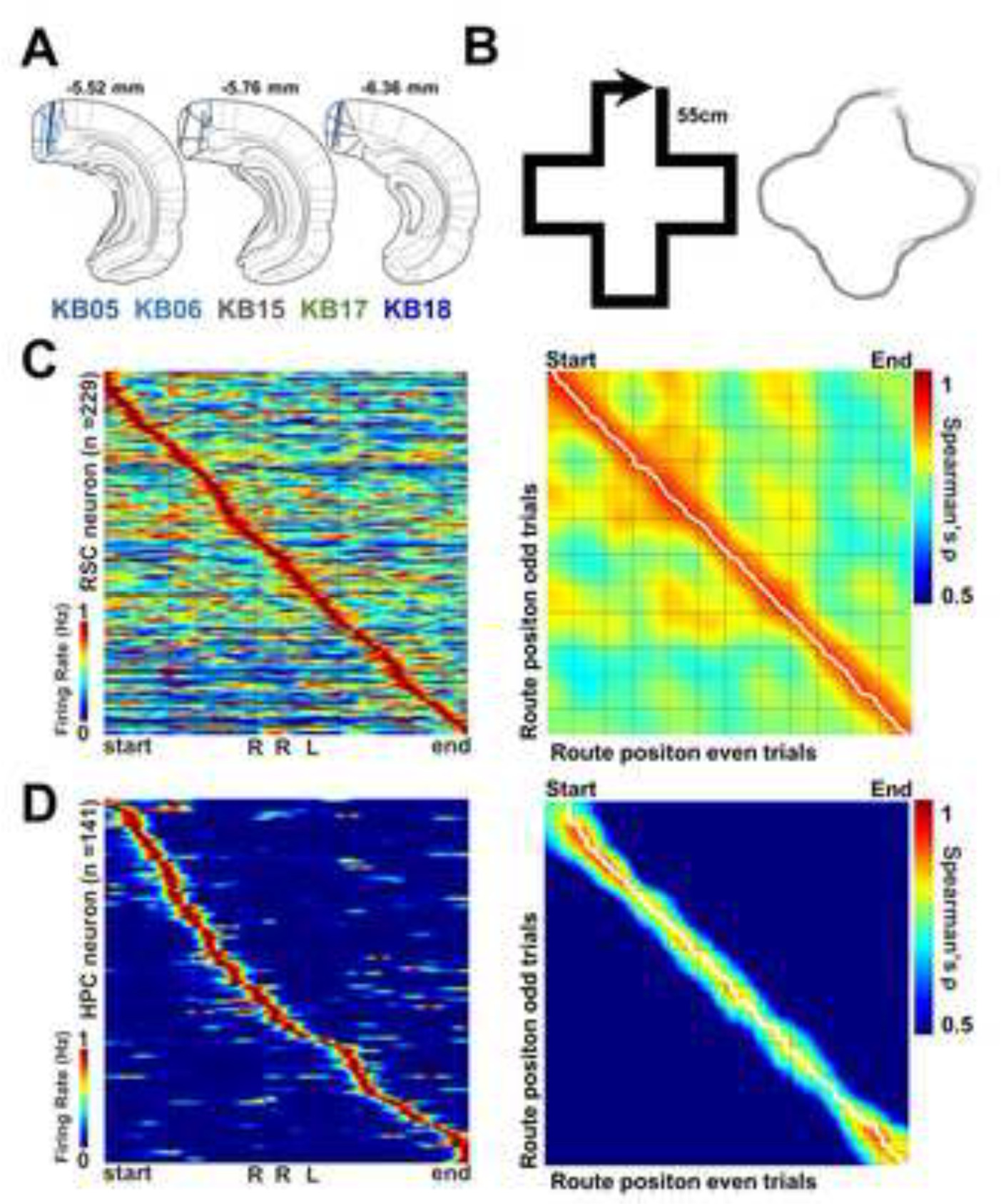
RSC encodes progression through a closed track with recurrent structure. **A.** Placement of electrode wires across all rats (n = 5) in caudal RSC collapsed across hemispheres. Lines indicate recording depths included for analysis. Line color indicates rat identity. **B.** Left, schematic of the ‘plus’ track. Track is closed, in that the animal starts and ends at the same location and is rewarded at the arrow point. Right, tracking of multiple ballistic traversals of an individual animal in a single session. **C.** Max normalized firing rate profiles across all RSC neurons (n = 229), sorted by position along track with maximal firing rate. High firing along the diagonal indicates that a unique population of neurons exhibited peaks in activation for every position along the track. Right, for the same population as on the left, a correlation matrix of population vector similarity computed from odd and even numbered trials. Strong correlation along the diagonal indicates that across non-overlapping trials, the maximal similarity between population vectors occurred at the same position through the route. White line, peak correlation for each row showing that the animals position along the track can be accurately decoded. Off diagonal red indicates repetition of activation patterns for different positions along the track and thus, potential presence of spatial periodicity. For both plots, dashed lines show position of left turns, non-dashed lines show position of right turns. **D.** Same as plots in C but for HPC pyramidal neurons (n = 141).

To determine whether RSC ensembles generate unique and reliable patterns for every route position, we split route traversals into odd and even trials, generated separate positional rate vectors for each, and created a correlation matrix which assessed the degree of similarity of ensemble patterns for each route position against all others (**Figure 1C, right**). The extent to which maximal similarity between odd and even trials lies along the upper-left to lower-right diagonal of such matrices provides a measure of the extent to which unique, reliable patterns are formed at each location^12^. Such ‘reconstruction’ of the animal’s progression along the route was accurate to 1.9 ± 2.7 centimeters based on RSC populations, a value which compared favorably to the accuracy seen for a population of 141 hippocampal CA1 ‘place cells’ recorded in 3 animals (**Figures 1C, 1D;** CA1 accuracy = 5.2 ± 6.7 centimeters).

Positional correlation patterns for RSC and CA1 neurons were distinct, however, in the off-diagonal correlations that reflect similarity between non-adjacent track locations. Unlike matrices for the CA1 ensemble, off-diagonal correlations for RSC ensembles exhibited repeating patterns across the track. The spatial distribution of such off-diagonal correlations suggests, in particular, the presence of a sub-population of RSC neurons having repeating patterns over quarter and/or half sections of the full route.

### RSC neurons exhibit spatial periodicity at full route and sub-route spatial scales

As stated, spatially periodic activation patterns in RSC ensembles could reflect concurrent encoding of route sub-spaces by individual neurons. To examine this question directly, we applied generalized linear models (GLMs) to determine whether individual RSC cells exhibit activity peaks that follow the recurrence of sub-spaces at multiple scales inherent to the spatial structure of the route.

The GLM analysis was designed to fit the positional firing rate profiles of individual neurons. We constructed multiple predictors (**Figure 2A**) composed of paired sin and cosine functions with different pairs having periods corresponding to full route or sub-route spaces. Each pair acts together as a single predictor having a distinct set of values for each position in the full route space or having repeating sets of values across same-sized sub-spaces. The latter are used to detect recurrence in firing patterns for analogous sub-spaces. We chose the predictors to reflect the logical fractionations of the space into twelfths, corresponding to the smallest track segments, through sixths, quarters, thirds, halves, and the full space of the track.

**Figure 2.**
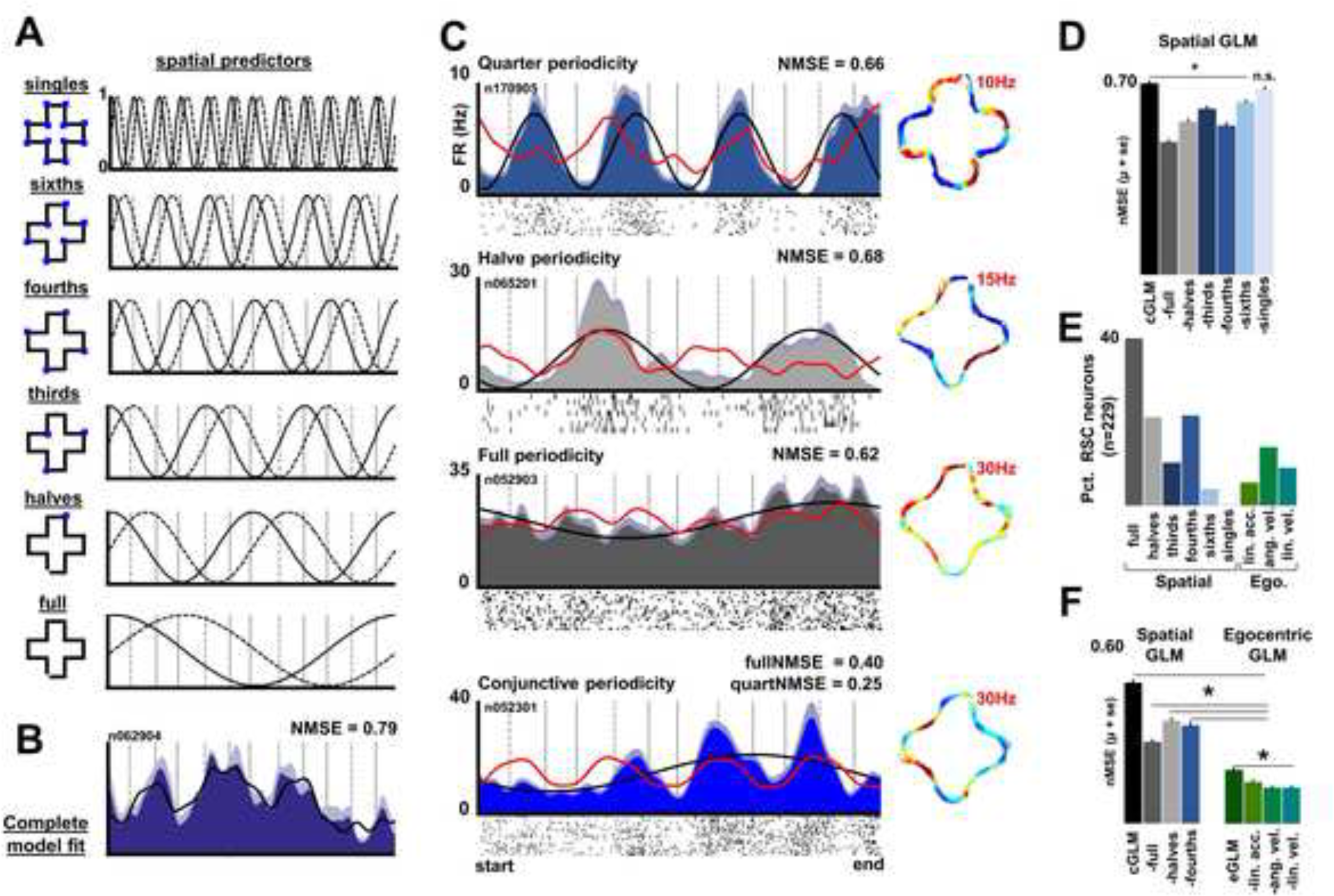
RSC neurons exhibit spatial periodicity. **A.** Left column, schematic of fragmentation of plus track into multiples of 1/12 using oscillating spatial predictors constructed from sine and cosine functions. Track space is split into twelfths, sixths, quarters, thirds, halves, and full fragments. Blue squares indicate the boundaries of fragmentations. For each fragmentation, cosine and sine functions were constructed that cycled at the scale of the full track space or for the corresponding spatial scale. Spatial predictors are shown in the plots to the right of the schematic for each fragmentation. **B.** Example of complete GLM (cGLM) model fit, using all spatial predictors, generated for the mean firing rate vector (± s.e.) for an individual RSC neuron. The black line is the complete GLM fit using all predictors and the fit between actual and predicted vectors is indicated above the plot (normalized mean squared error-NMSE, 1 is a perfect fit). **C.** Four mean firing rate profiles for individual RSC neurons showing different scales of spatial periodicity. For each plot, the model fit generated by the best individual spatial predictor (iGLM) is shown in black, and the NMSE is indicated above the plot. In red, the partial GLM (pGLM) when the best predictor is dropped from the model showing the impact of that spatial periodicity in accounting for the firing pattern of the neuron. Below each plot, spike trains for individual trials. To the right of each figure is the corresponding two dimensional ratemap with a max threshold indicated in red (zero firing rate is blue). **D.** Population statistics for model fits. Across all RSC neurons, the mean (± std. error) fit of the cGLM (black bar) compared to all pGLMs (gray and blue bars). All pGLMs had significantly lower fits on average with the exception of the pGLM that fragmented the plus into individual segments (‘singles’, Kruskal-Wallis, X^2^(6) = 250.5, *p* = 3.2 × 10^−51^, *post hoc* Bonferonni correction). **E.** For each neuron and trial, a cGLM and egocentric GLM (eGLM) was generated. All pGLMs (both spatial and egocentric) were then generated for that trial and NMSE was computed. This process was repeated across all trials for an individual neuron to generate a distribution of trial-by-trial pGLM values that were then significance tested against the same distribution for trial-by-trial cGLM values. Shown here, percentage of RSC neurons significantly impacted by each spatial fragmentation or egocentric-based movement variable. **F.** Spatial versus egocentric predictors. Left, spatial cGLM and spatial pGLMs using the three best spatial predictors from 2D. Right, complete eGLM and partial eGLMs constructed with linear velocity, acceleration, and angular velocity as predictors. All eGLM predictors significantly impact the complete eGLM. Spatial GLM models are significantly better at fitting RSC neural data than models generated with egocentric data as predictors.

To quantify model fits to the actual positional rate vectors, we used the normalized mean squared error (NMSE) between the rate vector of any given neuron and a model that used all of the spatial predictors (the ‘complete model’, or cGLM; **Figure 2B**). To assess the contribution of each spatial predictor to model accuracy, we then removed each spatial predictor in isolation. We refer to these as partial GLMs (pGLMs). The increment in NMSE for pGLMs versus cGLMs corresponds to the decrement in fit and measures the strength of the removed predictor in improving cGLM fits. To complement this approach to measuring the influence of the full route or sub-route spaces on the firing pattern of each neuron, we also calculated NMSEs for models generated using only individual spatial predictors (iGLMs).

**Figure 2C** shows the positional rate vectors of four individual RSC neurons with the best single-predictor fit (best iGLM model fit) overlaid in black and the pGLM model fit with that best single-predictor removed (overlay in red). Applying this same methodology to the full RSC population, the fit between the best individual predictor (based on iGLM models) was found to be 2.3 ± 2 times larger on average than to the second best, indicating that most RSC neuron firing patterns are dominated by a single full route or sub-space periodicity.

Similarly, examining the fit decrements in pGLMs relative to their associated cGLMs across the full RSC population, we found that all but one predictor significantly impacted the complete model fit (**Figure 2D**, Kruskal-Wallis, X^2^(6) = 250.5, *p* = 3.2 × 10^−51^, *post hoc* Bonferonni correction). The only exception to this was the ‘singles’ predictor corresponding to the single segment spatial scale.

To statistically test the impact of each full and sub-route periodicity on the firing of individual neurons, we calculated fit correlations for pGLMs and cGLMs for the positional rate vectors associated with individual trials. In this analysis, an assessment of each predictor’s contribution to strength of fit is calculated for each run through the full route. The distributions of fit values for pGLMs versus cGLMs are then compared. The percentage of RSC neurons significantly impacted by each predictor is shown in **Figure 2E** (individual neuron Kruskal-Wallis across trials, *post hoc* Bonferonni, p<0.05).

Consistent with the results of the aforementioned GLM approaches and with the structure of the correlation matrix given in **Figure 1C**, 63.7% of RSC neurons (n = 146/229) exhibited periodicity in firing according to at least one full-route or sub-route space. Specifically, 47.2% (n = 108/229) of the neurons were modulated by sub-routes as manifested by significant spatial periodicity at scales subordinate to the full route. Of RSC neurons that were sub-route modulated, the most prominent encoding fell at quarters (21%, n = 48/229) and halves (20.5%, n = 47/229). 38.9% (n = 89/229) of RSC cells had an activation pattern that cycled a single time across the full space of the trajectory. Example neurons with periodicity in firing rate patterns over quarter and half sections of the full route and with periodicity over the full route space are given in **Figure 3**. Populations of sub-route encoding neurons and those that exhibited ‘full’ oscillatory behavior were not mutually exclusive, as 48.3% of ‘full’ neurons (n = 43/89) were also conjunctively sensitive to one or more sub-routes. Of all neurons, 26.6% (n = 61/229) were conjunctively sensitive to multiple spatial scales simultaneously (**Figure 2C bottom, Supplemental Figure S2**). This latter finding illustrates a neural framework by which sub-route relationships can be hierarchically mapped to other sub-routes or the full trajectory.

**Figure 3.**
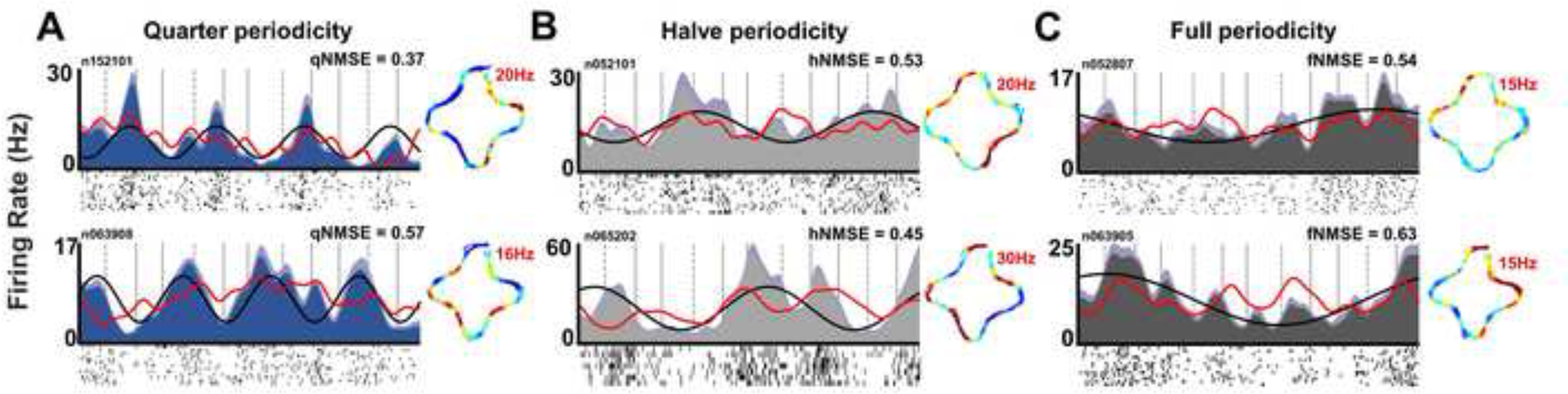
RSC neurons exhibit spatial periodicity biased to quarter, half, and full spatial scales. **A.** Mean linear firing rate vector (± s.e.) of two example RSC neurons exhibiting significant modulation that oscillates four times across the full route. For each plot, the black line is the iGLM fit using the best predictor. In red, cGLM fit with best predictor dropped showing decrement in fit when best periodicity is no longer included. Below, spike trains for individual trials showing reliability of firing patterns across track space. Right, corresponding 2D ratemap showing where the activation occurs in the full space of the room. Red number indicates peak firing rate in Hz for these plots (blue is zero firing). NMSE, normalized mean squared error of best model (prefixes - q = quarter, h = halve, f = full). **B** and **C**, same as **A**, but for RSC neurons exhibiting periodicity that oscillates across half of the track and the full track space, respectively.

### Quarter section periodicity is not accounted for by locomotor actions

RSC neurons are capable of generating reliable responses to specific actions such as left or right turns, though such responses are not correlated to variation in angular velocity on a trial-by-trial basis^5^. For the plus-shaped route utilized here, quarter sections of the full route represent the smallest sub-space over which such specific actions repeat. Consistent with this feature of the route, movement profiles were well described by spatial predictors that oscillated at the scale of quarters across the full trajectory (**Supplemental Figure S3**). For these reasons, it was important to determine whether some or all of the sub-space periodicity in firing, particularly at the level of quarter sections, could be explained by correlation of firing rates to specific locomotor variables such as linear and angular velocity and linear acceleration. Full-route and half-route periodicity in firing is unlikely to be explained by locomotor action correlates because those correlates repeat over quarter sections. Nevertheless, we considered locomotor variables for their potential contribution to the firing of all RSC neurons and for all full and sub-route spaces.

To do so, we generated GLM fits for each neuron’s positional rate vector using predictors based on the actual linear and angular velocities and linear acceleration profiles observed during track-running. We refer to these as eGLMs with ‘e’ itself denoting the egocentric nature of the movement variables. Overall, eGLM fits indicated that such movement variable predictors produced poorer fits than the previously-described spatial predictors, even when matched for total number of predictors in the model (eGLM; **Figure 2F, Supplemental Figure S3,** Kruskal-Wallis, X^2^(7) = 771.2, *p* = 3.07 < 10^−162^, *post hoc* Bonferonni correction). Thus, moment-to-moment fluctuations in RSC firing are not well described by moment-to-moment fluctuations in movement variables.

The proportion of individual RSC neurons significantly impacted by fluctuations in the aforementioned movement variables was also assessed with a trial-by-trial eGLM analysis. Here, firing rate predictors for each trial were constructed using the corresponding trial’s linear acceleration, velocity, and angular velocity vectors. Only 5.2% (n = 12/229) and 8.7% (n = 20/229) of RSC neurons were significantly impacted by linear acceleration and velocity, respectively. 13.5% (n = 31/229) of trial-by-trial eGLM model fits to RSC neurons were significantly decremented by the loss of angular velocity predictors (**Figure 2E**).

Finally, despite the overlay of spatial and movement variable periodicity at the level of route quarter sections, just 43.7% (n = 21/48) of neurons that exhibited quarter-scale spatial periodicity also exhibited significant sensitivity to any of the movement variables as assessed with this trial-based eGLM approach. These findings indicate that the presence of quarter periodicity is, in general, not directly reflective of recurrence in movement variables.

### Spatial firing of RSC neurons is anchored to route and allocentric spaces

Sub-route encoding in the form of periodic activation could potentially be explained by factors other than spatial relationships within route or allocentric space. As observed in other structures, RSC activation patterns could reflect path integration anchored to locations associated with actions, trajectory start points, or trajectory endpoints associated with reward. For example, grid cells of the medial entorhinal cortex (mEC) exhibit repeating activation patterns during track running and such patterns can be anchored to the start points of track segments^13,14^ or to reward locations^15^. Similar findings have been reported in the hippocampus and striatum, where activation of neurons can reflect the animal’s future trajectory to a reward site^16–18^. A final possible explanation for RSC periodicity could be firing related to passage of time during constant locomotion as observed in the HPC^19^ and mEC^20^.

To test whether periodic activation of RSC neurons was related to distance from starting or ending location, time from start or end of trial, or a combination of route and allocentric spatial position, we trained two rats to run randomly interspersed half and full routes within the same plus track environment (**Figure 4A**). The half trajectory was positioned over the middle portion of the full route which displaced the start point in allocentric space. Rats were required to utilize distal cue information to identify the route type of each trial, and without cuing, either stop or continue through the position on the plus track associated with the endpoint of the half route. 101 RSC neurons were recorded under these conditions.

**Figure 4.**
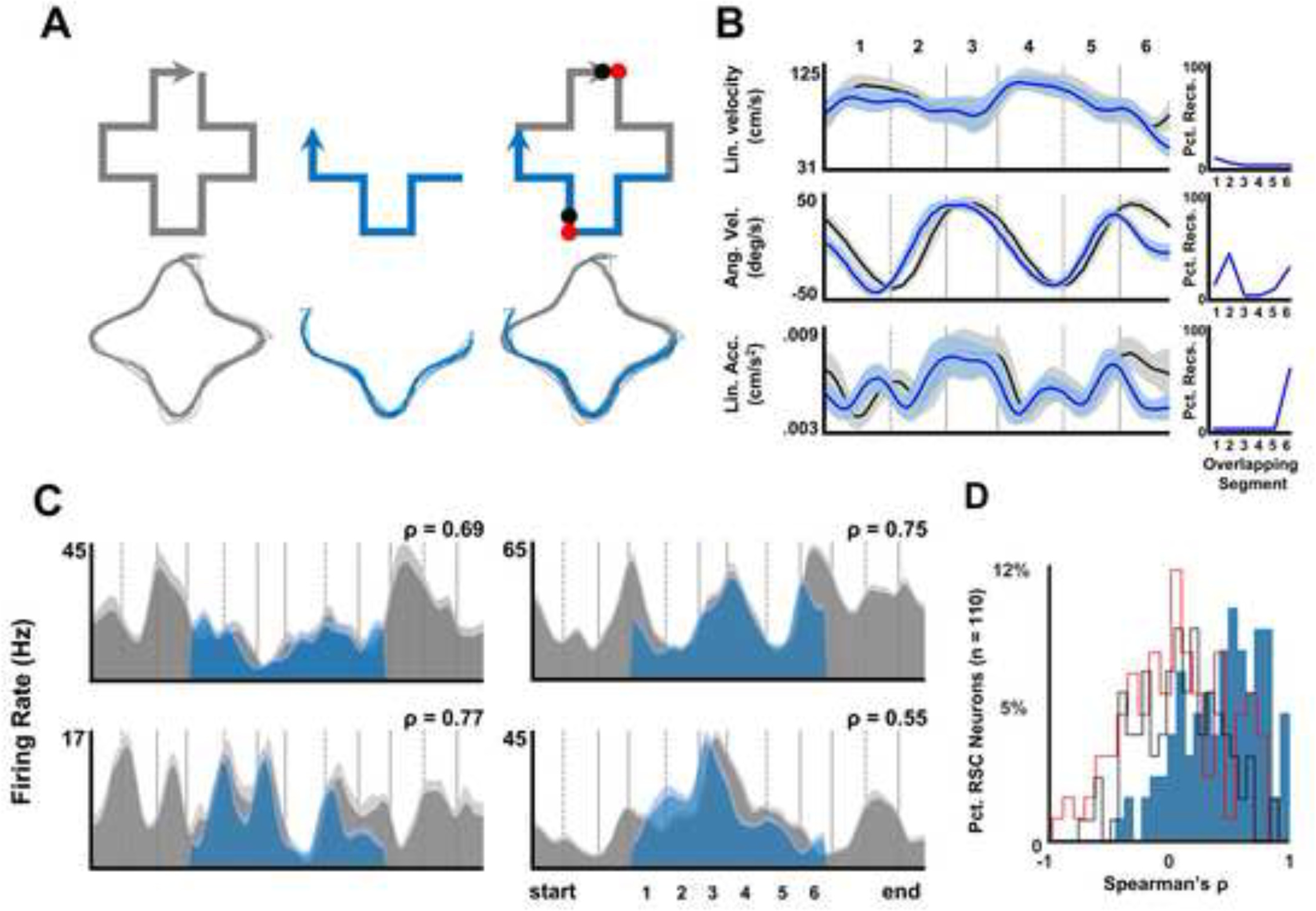
RSC activation patterns are anchored to allocentric space. **A.** Rats (n =2) were trained to run randomly interspersed full and half traversal routes, starting and ending at two different locations within the same plus track. Top row left, schematic of full route trajectories. Top row middle, schematic of half route. Top row right, schematic of route overlap. Red circles depict boundaries of first half of full route. Black circles depict boundaries of last half of full route. Bottom, example tracking data from a single animal during a single recording session showing ballistic traversals for both routes. **B.** Mean (± s.d.) of linear velocity, rectified angular velocity, and linear acceleration movement variables across all rats and recording sessions for the overlapping portions of the track for each route. Blue and black vectors represent half and full route traversals, respectively. Overlapping segments are numbered. To the right of each figure is the percentage of recordings (n=40) with statistically significant differences between half and full routes for each segment (Kruskal-wallis test, post hoc Tukey-Kramer, p<0.05). Most significant differences occur on the last (6) segments, where the animal is stopping or continuing through the segment depending on current route. C. Mean (± s.e.) firing rate profiles for 4 individual RSC neurons during traversals of half and full routes (blue and black profiles). For each neuron, the correlation between activation profiles in overlapping space are shown at the top right. Despite complex firing rates, a large portion of neurons exhibited strong spatial correlations in firing rate across the two routes, indicating that the activation pattern was not anchored to the distance from the beginning of the route, nor to time. Instead, spatial firing patterns are anchored to the allocentric position in the environment. Segments are numbered to match **4B. D.** Distribution of half to full correlations for all neurons recorded under these conditions (n =110). The distribution for overlapping segments (blue bars) is strongly biased to positive correlation values. In red, distribution of half correlations to the first half of full runs rather than overlapping track segments. In black, distribution of half correlations to the last half of full runs rather than overlapping track segments.

Locomotor variables for the overlapping track segments of full route and half route journeys were remarkably similar (**Figure 4B**). Within each recording, movement through most shared segments did not statistically diverge between the two routes (**Figure 4B, right;** n = 40 recordings, Kruskal-Wallis, *post hoc* Tukey-Kramer, p > 0.05). The final segment did have statistically different linear acceleration in a majority of recording sessions (60%, n = 24/40) as the animal either stopped or continued through this segment depending on trial type (Kruskal-Wallis, *post hoc* Tukey Kramer, p<0.05). However, the overall similarity of movement indicated that any RSC firing pattern differences found between full and half routes were not conflated with differences associated with the rat’s movement.

We next examined the relationship between RSC firing rate profiles between half and full trajectories. If RSC activation was related to path integration as a function of either distance or time from the start or end point, we expected the pattern on the half route to closely match the first half (**Figure 4A, top right, track locations between the red circles**) or last half of the full route (**Figure 4A, top right, track locations between the black circles**), respectively. In contrast, if activation was anchored to space, we expected the overlapping portions of the half and full routes to exhibit strongly correlated profiles. This latter hypothesis, in which activation was anchored to the external environment, was the overwhelming trend. **Figure 4C** depicts four example RSC neuron firing patterns across full and half trajectories. Correlation values for the four neurons reveal strong similarity in patterning between the overlapping portions of the half and full route traversals. Further, correlations between half and full routes in overlapping space were statistically greater than those computed between the half route and both the first half and last half of full route traversals (**Figure 4D**, n = 101, μOverlap = 0.45, μFirstHalf = 0.09, Wilcoxon rank-sum test, z = 6.74, p = 1. 60 x 10^−11^; μSecondHalf = 0.08, Wilcoxon rank-sum test, z = 7.26, p = 3.80 × 10^−13^). As such, we conclude that the RSC firing profiles described here are spatially anchored and not the product of distance or temporal integration from the start or end of trials.

### Spatial periodicity in RSC firing does not strictly demand recurrence in action sequences

A striking feature of the spatially-anchored periodic activation found in RSC was the apparent bias to quarter and half sub-routes. These sub-route representations segmented the full trajectory into equivalent action sequences that collectively construct the full route. Although the eGLM results demonstrated that the spatial periodicity is not well explained by a direct relationship between firing rate and action correlates, the question remained whether this form of spatial representation would persist in environments that did not have recurrent structure that could explicitly define boundaries of route fragments. To examine this question, rats (n = 3) were trained to run a ring shaped track in the clockwise direction. The ring track lacked action sequences and animals held consistent angular and linear velocity profiles for the majority of ring traversals (**Supplemental Figure S4**). 90 RSC neurons were recorded under these conditions.

As observed in RSC activation patterns during plus traversals, the spatial cGLM analysis revealed sub-route representations in the form of spatially periodic activation patterns during ring track running. Most spatial predictors significantly impacted the complete model fit, the exceptions being those that segmented the ring track into sixths and twelfths (**Figure 5A, left,** X^2^(6) = 180.2, *p* = 3 × 10^−36^, *post hoc* Bonferonni correction). 86.7% (n=78/90) of RSC neurons were significantly modulated by at least one oscillating spatial predictor when analyzed across trials (**Figure 5B**, gray and blue bars). Surprisingly, sensitivity to a particular sub-route did not generalize between the plus and ring, as relatively few of the neurons recorded on both tracks (n = 78/90) were statistically sensitive to the same predictor in the two route running conditions (**Figure 5B**, black bars). Proportions of neurons whose firing patterns were significantly predicted by linear acceleration, speed, and angular velocity (eGLM) were low (<10%) and less than proportions observed on the plus (27.5%, **Figure 5B**, green bars). For neurons that exhibited significant modulation by full, half, or quarter spatial predictors, individual predictor model fits were as good on the ring as those for neurons exhibiting the same scales of spatial periodicity on the plus (**Figure 5C**, ‘Full’, μRing = 0.41, n = 63, μPlus = 0.35, n = 89, Wilcoxon rank-sum test, z = 1.32, p = 0.19; ‘Half’, μRing = 0.27, n = 40, μPlus = 0.29, n = 47, z = −0.95, p = 0.34; ‘Quarter’, μRing = 0.36, n = 12, | μPlus = 0.33, n = 48, z = 0.49, p = 0.62).

**Figure 5.**
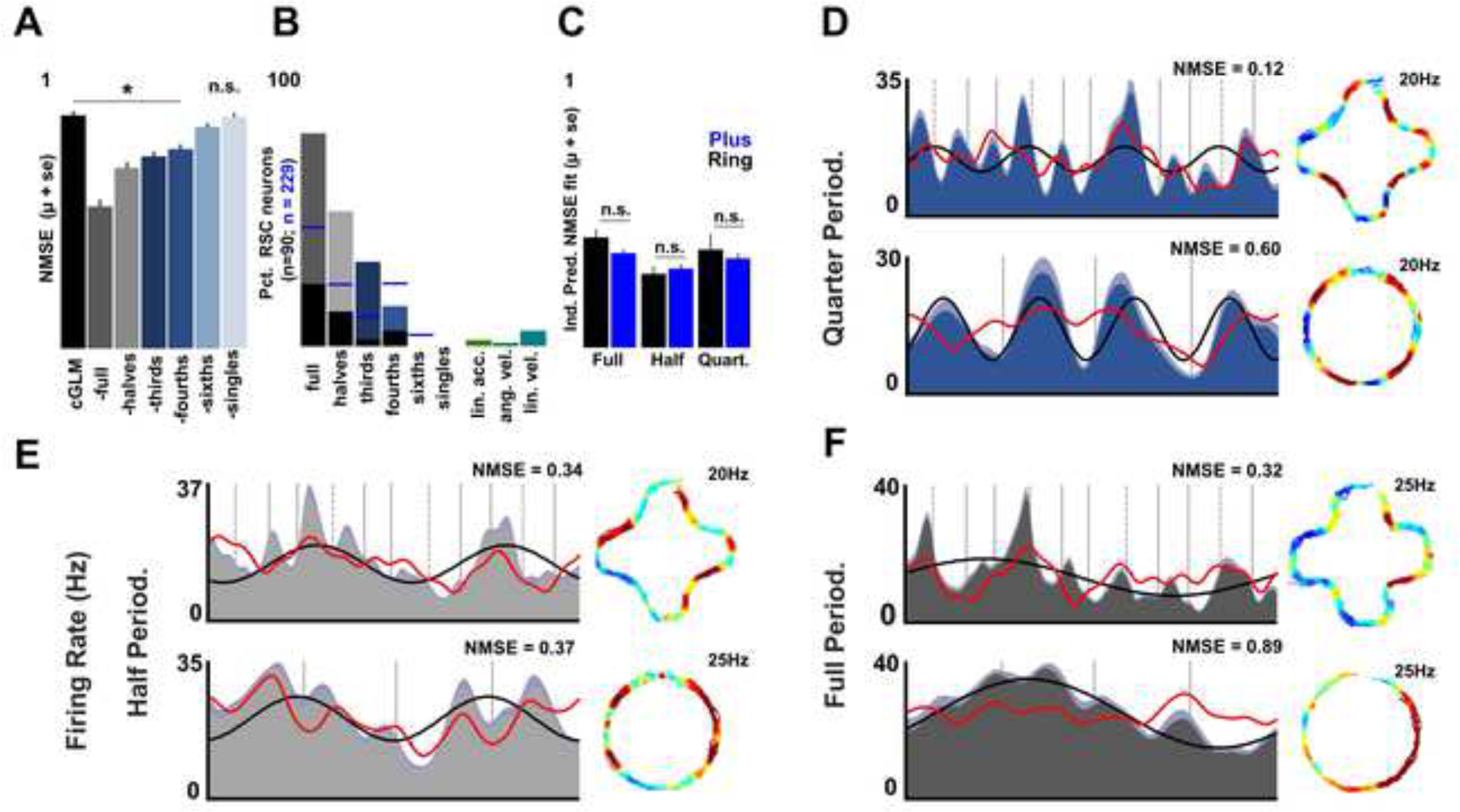
RSC neurons exhibit spatially periodic activation patterns on a ring shaped track. **A.** Across all RSC neurons recorded on the ring track, the mean (± s.e.) NMSE of cGLMs and pGLMs. With the exception of ‘single’ and ‘sixths’ spatial periodicity, all other spatial predictors significantly impacted the complete model (X^2^(6) = 180.2, *p* = 3 × 10^−36^, post hoc Bonferonni correction). **B.** For each neuron and trial, a cGLM and eGLM was generated. All pGLMs (both spatial and egocentric) were then generated for that trial and NMSE was computed. This process was repeated across all trials for an individual neuron to generate a distribution of trial-by-trial pGLM values that were then significance tested against the same distribution for trial-by-trial cGLM values. Shown here, percentage of RSC neurons significantly impacted by each spatial fragmentation or egocentric-based movement variable on the ring track. Black bars, percentage of all neurons recorded on the ring (n = 90), that were significantly modulated by the same spatial predictor for both ring and plus track sessions. Blue lines, percentage of RSC neurons recorded during plus track running that were significantly modulated by each spatial predictor (same as bars in **Figure 2F**). **C.** Mean (± s.e.) of fit to individual predictors for all neurons that exhibited significant modulation by full, halve, or quarter spatial predictors in either track running condition (blue, plus; black, ring). Fit was not statistically different between track types, indicating that the spatial periodicity was equally strong across conditions **D.** Mean activation profiles for a single rSc neuron that exhibited significant modulation at the quarter scale during ring traversals (bottom, ring; top, plus). Right, corresponding 2D ratemaps. **E and F.** Same as **4D**, but for two RSC neurons that exhibited significant modulation at half and full scales during ring track running, respectively.

However, the proportions of RSC neurons that were significantly modulated by individual predictors differed significantly for the plus and ring tracks, with spatial periodicity on the ring being biased to larger scale oscillations (**Figure 5B;** Chi-squared test, X^2^(3) = 80.83, *p* = 2.03 × 10^−17^). **Figures 5D-F** show the activation profiles, for both ring (bottom) and plus (top) sessions, of RSC neurons that exhibited significant spatially periodicity on the ring. Although RSC neurons exhibited quarter periodicity on the ring, the frequency of quarter sub-route encoding (13.3%) was diminished compared to the plus track (21%, **Figure 5D**). In contrast, the proportion of neurons that were modulated at the scale of half of the route was higher for the ring (44.4%) than the plus (20.5%, **Figure 5E**). Finally, 70% (n =63/90) of RSC neurons exhibited periodic patterns on the track that oscillated at the scale of the full ring traversal (**Figure 5F**), in contrast to 38.9% during plus track traversals. These differences suggest that, although sub-route extraction is an inherent function of the RSC network, the proportion of neurons sensitive to specific sub-routes is adaptable and reflects the structure of the current environment.

### Full route periodicity in RSC firing yields a map of distances between all route locations

Spatial periodicity of the form observed in RSC provides a neural mechanism for extracting analogous positions within repeating sub-trajectories. This encoding is manifested as a distinct firing rate code and pattern across locations within a sub-route, that then repeats for each iteration of the sub-route within the full trajectory. Many RSC neurons exhibited sensitivity to repeating sub-routes in the form of periodic activation patterns, yet an equally large number exhibited oscillatory patterns that scaled the full space of the route. The spatial representation for these ‘full’ neurons is unique relative to other spatial mappings currently reported. Here, activation is spatially reliable, continuous (rarely exhibits zero-firing), cyclic, and exhibits symmetrical firing activation across two complete halves of the track. This symmetry is in stark contrast to that observed for HPC place cells, as the scale of symmetry covers the entire route space, not just positions within a localized firing field surrounded by zero-firing regions (**Figure 6A-B**).

**Figure 6.**
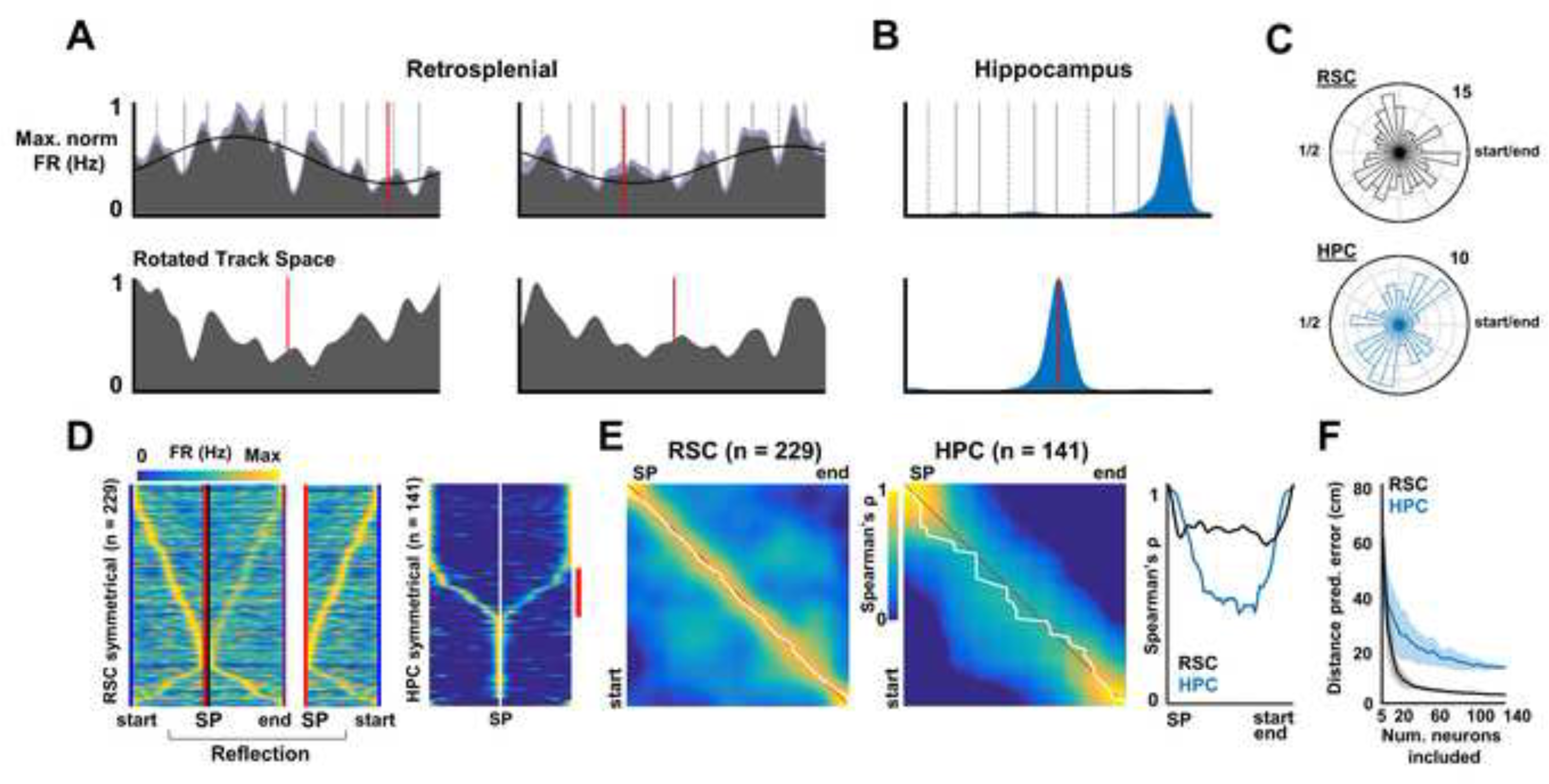
RSC spatial tuning yields a metric for distance. **A.** Top row, mean firing rate profiles for two RSC neurons on the plus track. Red line depicts ‘symmetry point’ where rotation of the vector yields maximal symmetry. Second row, rotated versions of top row plots that yield greatest reflective symmetry. Red dashed line shows that the symmetry point is now in the center of the firing rate vector. **B.** Same plots as in 5A, but for a HPC place cell. **C.** Distribution of symmetry points along the plus track. **D.** Left, max-normalized firing rate profiles for all RSC neurons aligned to points of symmetry (red line) and sorted by position of peak activity. The left half of this matrix, from the adjacent vertical blue line to the red line in the left panel, was extracted and reflected (middle panel, notice flipped positions of red and blue lines). Population vectors from the reflection of the first half were then correlated with population vectors taken from the second half of the left matrix (from the black to the purple line). Right, symmetrical max-normalized firing rate profiles for all HPC neurons. Red bar to right of HPC symmetrical matrix, population of HPC neurons with multiple place fields. **E.** Distance correlation matrices and distance error estimation. Left and middle, distance correlation matrices reflect the similarity of RSC or HPC population activity as a function of distance in opposite directions from the point of symmetry. Strong correlations along the diagonal, from upper left to lower right, demonstrate strong similarity for positions equidistant to the symmetry point. A distance estimate of the animal’s position from the symmetry point is computed by finding the maximal correlation in each row. This value is depicted by the white line. A perfect reconstruction of the animal’s distance would run diagonally from upper left to lower right, as shown by the red line. The error in distance estimate is computed by finding the absolute value of the difference between each point along the white line and the corresponding point on the red line. Right, correlation values along the diagonal for RSC and HPC distance correlation matrices. **F.** Mean (± s.d.) distance reconstruction error of RSC neurons and spatially-responsive HPC neurons (place cells) as a function of the number of neurons from each region included in the analysis.

This novel form of spatial encoding, in turn, generates a novel form of spatial information. Specifically, symmetrical firing patterns anchored to route space provide information about the animal’s distance from a fixed point. To assess the presence of a distance code, all RSC firing rate profiles collected on the plus track were rotated to find the best point of symmetry (**Figure 6A, bottom**). Such symmetry points (SP) were those that reduced the error (again, NMSE) when the surrounding halves of the track were compared to each other. Symmetry points were uniformly distributed across the route space for RSC and HPC (**Figure 6C,** Rayleigh test, RSC, n = 229, z = 0.73, p = 0.48; HPC, n = 141, z = 1.01, p = 0.37). Thus, it is likely that large sub-populations of neurons (including RSC neurons not recorded) have anchor points at each track location and we applied this assumption to our next set of analyses.

Examination of rotated and SP-aligned firing rate profiles revealed the presence of dual firing peaks at positions approximately equidistant from the SP (**Figure 6D, left panel**). This finding indicated that there was strong population vector similarity for track positions located at similar distances from the SP. For HPC neurons, the same process revealed symmetry but the portions of the rotated track space between peaks were generally associated with a lack of firing (**Figure 6D, right panel**).

To test whether the animal’s distance could be accurately predicted as a function of distance from the SP, we began by examining the complete RSC and HPC ensembles. Vectors for all RSC neurons were halved at the point of symmetry and the first half of the firing rate profiles were reflected as shown in **Figure 6D** (data from blue to red vertical bars is reflected from left to right and depicted in the middle panel, now from red to blue vertical bars). Distance reconstruction was computed by correlating the RSC population vectors for each position moving away from the SP with each reflected position moving towards the SP (**Figure 6D**). The correlation matrix produced from this process is shown in **Figure 6E** for the RSC and HPC. Strong correlation along the diagonal reflects the presence of a similar population vector for positions equidistant, but occurring on opposing sides, of the SP. The animal’s predicted distance from the SP was simply taken as the point associated with maximal correlation for each row (shown in white in **Figure 6E**) and the error in distance reconstruction is the absolute difference between each position on the white line and a perfect distance prediction line (shown in red in **Figure 6E**). The mean distance reconstruction error for the RSC was 2.09 centimeters and significantly lower than that observed in HPC (μ_HPC_ = 12.97 cm, Wilcoxon rank-sum test, n = 72, z = 6.96, p = 3.44 × 10^−12^). Any semblance of HPC distance representation was likely afforded by a sub-population of neurons that exhibited multiple, distinct, firing fields across the track (highlighted by the vertical red bar running to the right of the right panel in **Figure 6D**). By and large, HPC distance correlations were much lower and generally less reliable than those observed in RSC (**Figure 6E, right**).

Although the difference in distance encoding between RSC and HPC was striking, the distance reconstruction analysis is sensitive to the absolute number of neurons included in the corresponding population vectors. Accordingly, we iteratively repeated the distance estimation process, each time utilizing a matched, randomly-chosen sub-sample of RSC or HPC neurons. For each quantity of randomly sub-sampled neurons, the mean error in predicted distance was computed 100 times. The average distance reconstruction error was significantly lower in RSC than HPC, beginning when only 10 neurons were utilized to compose the population vectors used in the distance estimation (**Figure 6F,** Wilcoxon rank-sum test, n = 200, z = 5.55, *p* = 2.9×10^−8^).

Symmetrical RSC ensembles were significantly better than HPC at predicting the animals distance from the SP, even as up to 140 neurons (equivalent to the total recorded HPC population) were added to the population vectors (two-way ANOVA between brain regions with *post hoc* Bonferroni test, F(1) = 5780, *p* = 0). For both populations, mean error in distance reconstruction across 100 iterations began to asymptote when approximately 20 neurons were utilized to compose the population vectors (**Figure 6F**). Collectively, these findings demonstrate a unique capability of RSC to encode distance from uniformly distributed points of symmetry. This property of RSC ensembles is afforded by the presence of RSC neurons that exhibit continuous, cyclic, activation profiles across the full route space, a spatial representation not observed in the HPC.

## Discussion

The current experiments and analyses addressed RSC spatially-specific firing patterns in the context of a defined route having recurrence at multiple spatial scales. RSC ensembles, like CA1 ensembles, exhibited unique firing patterns for every location of the plus-shaped route consistent with recently published work^5^. However, the results also reveal unique properties of spatially related activation within RSC that contrast to firing patterns of dorsal CA1 neurons. First, sub-populations of RSC neurons exhibited recurrent firing patterns that effectively define analogous route sub-spaces at multiple spatial scales. Examination of neurons during traversal of circular tracks indicates that such decomposition of routes into sub-spaces may be a general property of RSC, one not absolutely dependent on recurrence in path structure nor action sequence. Second, we show that the spatially-specific, but continuously-firing nature of a large RSC sub-population yields a mapping of distance travelled and distance between any given track position and all other track positions. Together, the results implicate RSC in the extraction and encoding of the multiple spatial relationships inherent to the structure of complex paths.

We report a subset of RSC neurons that encode position within sub-routes of complex trajectories. More than half of RSC neurons recorded under the current conditions exhibited activation profiles that cycled multiple times across the full route. Because of this, similar firing rate patterns among the full RSC ensemble are observed for multiple route locations that have analogy with respect to position within sub-routes. At the single neuron level, such sub-space encoding was quantified by application of a GLM approach that assessed the degree to which model fits to trial-by-trial positional rate vectors were improved by predictors having different spatial frequencies. At the population level, recurrence in subspace patterns was found in correlation matrices mapping the full RSC pattern seen at each position to all other positions. Here, significant correlation peaks were observed for analogous route positions with spacing at the scale of route quarter sections. This result is consistent with the predominance of quarter-scale and half-scale firing pattern recurrence seen among individual RSC neurons.

Periodic activation patterns of this nature could be utilized to assess position within any individual route fragment, but, moreover, to also assess the position of fragments relative to each other and to the space of the full route. Two features of RSC spatially-specific firing are critical in this respect. First, consistent with our previous work^5^, RSC ensembles generated reliable and distinct population vectors for every position on the plus track. Despite the presence of strong correlations reflecting similarity among analogous positions of route subspaces, the strongest ensemble correlation values for odd versus even trials were for the same full route positions. Second, we also report the presence of RSC neurons that simultaneously encode multiple sub-spaces, via conjunctive periodicity. Notably, for a large number of such neurons, periodicity was simultaneously observed for the full route (single-cycle periodicity) and at scales subordinate to the full route space (e.g., quarter-section periodicity). These responses could potentially reflect neural mechanisms underlying hierarchical encoding and organization of spatial relationships^21–23^.

The periodic activation patterns observed most frequently segmented the full trajectory into quarters or halves. For all animals, traversals along the track also produced recurrent patterns in behavioral variables such as angular and linear velocity. Such recurrence was particularly obvious for quarter route subspaces. Thus, it was possible that some subspace encoding could be epiphenomenal to the animal’s behavior during navigation. Upon further consideration, several pieces of evidence are in conflict with this hypothesis. First, behavioral measures such as linear and angular velocity oscillate at the level of route quarter sections, but not route half sections. Thus, neurons with half-route periodicity exhibit firing rates that are different over the two quarter sections that compose each half section. Firing patterns for these neurons are dissociated from the patterns of linear and angular velocity. Second, a GLM analysis using behavioral predictors (linear and angular velocity and linear acceleration) produced significantly poorer fits to RSC firing rates than models constructed using spatial predictors. Finally, many RSC neurons exhibited strikingly clear sub-route activation patterns when the animal traversed a ring shaped track that lacked repeating action sequences. Sub-space patterning for the circle and plus-shaped tracks were often unrelated, demonstrating that subspace patterning on the circle was not driven by memory for the plus-shaped route structure.

Aside from demonstrating that periodicity in RSC spatial firing is not a simple product of motor behavior, the finding of periodic firing on the ring track suggests a more general tendency of the network to fragment spaces into subcomponents. Although the fitness of the spatial GLM was statistically the same across plus and ring tracks, the proportions of neurons that had oscillations at any given spatial scale shifted between the two routes. Thus, although RSC spatial fragmentation can exist without explicit sub-region demarcations (e.g. action sequences or local views of the track), the system can potentially be entrained to detect logical parsing of the environment when these cues are available.

The observed RSC sub-route representations should be considered with respect to novel forms of spatial representation recently reported in RSC. Specifically, cyclic activation patterns during route running on tracks having recurrence in shape could potentially reflect the position of the animal relative to local features of the track. For instance, an RSC neuron with quarter periodicity may fire on each straight track segment for which there are two turns visible to the right of the animal (see **Figure 2C, top**). Such an interpretation is consistent with recent fMRI work in humans demonstrating that multivoxel patterns in RSC are most similar for virtual locations that share the same locally-referenced position and heading direction^24^. A related phenomenon has recently been reported in free-foraging rats where a sub-population of dysgranular RSC neurons exhibited directional tuning referenced to the local visual cues and/or local boundary geometry in a multi-compartment environment^25^. Thus, the observation of subspace encoding in the present work is consistent with the idea that RSC neurons are sensitive to the proximal sensory features that define local spaces. Here we show that responses to local features can, in turn, be related to their positioning within the larger framework of the full environment. This response property, wherein RSC neurons are conjunctively sensitive to global position and proximal environmental features, has recently been proposed to anchor head direction representations to local sensory inputs^26^. Given the presence of direct RSC efferents to mEC, such dual encoding could also conceivably mediate transitions between local compartment and larger environment encoding of position as seen for mEC grid cells in multicompartment environments^27^.

Recurrence in spatial firing patterns has also been reported in several other structures, most notably in the form of grid cells of the mEC or repeating place fields of the HPC^27–40^. However, the current findings differ from previous work in several important ways. First, for nearly all cases in which CA1 and/or mEC neurons exhibit pattern recurrence, it is driven by exposure of the animal to identical or nearly identical proximal and distal landmark configurations. Here we find that RSC neurons exhibit pattern recurrence along routes that are visually open at all times to the changing views of the full recording environment and the distal cues at its boundaries. CA1 hippocampal neurons did not exhibit pattern recurrence in the same animals, indicating their ability to discriminate analogous route positions despite the aforementioned recurrences in local views of the track. Second, pattern recurrence in RSC neurons was observed on the circular track, the traversal of which yields identical views of the local environment. Third, RSC populations were found to conjunctively encode the full track space and multiple analogous track subspaces. Thus, considering the present findings in the context of prior work, it appears likely that RSC may be suited to generate recurrent patterning in the absence of such patterning within the HPC and mEC. It follows, then, that pattern recurrence in CA1 and mEC may depend on RSC projections that reach the HPC^41^ and mEC^42^ via direct and indirect routes. Nevertheless, confirmation of this awaits consideration of firing patterns of the subiculum which provides the most extensive input to the RSC and which is known to exhibit a higher degree of cross-environment generalization relative to CA1^43–46^.

Another structure in which sub-route representations have been found is the PPC^11^. PPC neurons exhibit repeating patterns along segments of nested routes that share similar actions and heading sequences, but which differ in allocentric position. RSC sub-route encoding diverges from the form found in PPC in two major ways. First, RSC can detect sub-route components that do not share the same heading orientation whereas PPC sub-route activation patterns were weakly correlated across different heading directions^11^. Notably, in most cases, the dependence of pattern recurrence on commonality in heading direction is also true for the aforementioned sub-space representations found in HPC^40^ or mEC^13^. A second key difference between periodic RSC patterns and those found in PPC, mEC, or HPC, is the larger spatial scale of the excitatory portion of the field, which, in conjunction with the cyclic nature of the response enables sub-route encoding and yields the metric for distance that we discuss next.

The current work reveals that ensembles of RSC neurons with periodic activation patterns collectively yield an ensemble code not only for distance from the animal’s current location (effectively 0 cm away), but also for distance to all other locations within the route. The existence of a RSC distance code is consistent with detriments to path integration following damage to the region^47,48^. Further, recent fMRI work has demonstrated strong RSC correlates with virtual distance integration^49–51^.

The RSC distance metric relies on activation profiles that wax and wane across the full track space. Critically, such activation profiles rarely have sections along the entire track where no firing occurs. Because of this and as a consequence of their sinusoidal shape, the spatial firing rate profiles for individual neurons possess a track location, here called the symmetry point, which segments the track into two mirrored, symmetrical halves. We demonstrate that RSC neurons possessing this property will exhibit highly similar firing rates for all locations approximately equidistant to the symmetry point location. This unusual property of RSC neurons and ensemble patterns stands in contrast to the observed firing patterns for individual CA1 neurons for which most track positions yield zero or near-zero firing. As a consequence, dorsal CA1 ensembles map distance only over spaces approximately the size of the average place field. Thus, at present, it appears that RSC is unique in simultaneously generating a mapping of position within path sub-spaces, current position in the full path space, and an ensemble code for distance between the current path position and all others. Of course, each of these properties may prove to exist in other brain sub-regions. In particular, it remains to be determined to what extent ventral CA1 populations, known to generate place fields of extensive length^52^ can effectively map distance.

Prior work demonstrates that RSC is also home to a small population of ‘head-direction’ neurons with precise tuning for the animal’s current head orientation relative to the larger environment^5,6,25^. Thus, it is plausible that the RSC distance encoding could be the integrated product of some combination of mEC grid cell, RSC head direction cell, HPC place cell signals, and RSC encoding of position within path sub-spaces. Consequently, we identify RSC as a spatial processing hub capable of highly unique computations relevant to efficient spatial representation, encoding of path structure, localization, and navigational problem-solving.

## Methods

### Subjects

Male Long-Evans rats (n = 5) served as behavioral subjects and were housed individually and kept on a 12-h light/dark cycle. Animals were habituated to the colony room and handled daily for a period of 1-2 weeks prior to shaping. Rats were food restricted to approximately 85-90% of their free-fed weight. Water was available continuously. All experimental protocols adhered to AALAC guidelines and were approved by IACUC and the UCSD Animal Care Program.

### Behavior

All animals were trained to traverse a plus-shaped track for reward. The track edges were 1 cm in height, which allowed the animal an unobstructed view of the full environment which had fixed distal cues across recording days. Animals ran the plus track in the clockwise direction.

Two rats were trained to run half traversals in addition to the full traversals that all rats ran. These same rats also performed track running on a W-shaped track in work that has been published^5^. On half route trials the rat started at the quarter point (of the full route) and stopped a quarter from the endpoint. The animal was required to stop at the correct endpoint on its own volition, without cuing from the experimenter. If the animal stopped correctly, a reward (1/4 honey nut cheerio) was placed on the track near the animal for consumption. Otherwise no reward was given. Half trials were randomly interspersed during all plus track sessions. For both full and half route traversals, animals were picked up between trials and carried in random trajectories to the trial initiation point.

A subset of rats (n=3) were also trained to traverse a ring-shaped track, in addition to the plus-shaped track. The ring track was centered on the same allocentric position as the plus-shaped track (relative to the recording room). Track edges on the ring were 1 cm in height. Ordering of exposure to the two tracks was counterbalanced across days to account for any potential sequence effects. Animals were trained to traverse the ring track in both directions, however clockwise data was solely analyzed as it allowed a more direct comparison to plus traversals given prominent head direction inputs to the RSC and observed directionality of spatial firing on linear environments. Fixed spatial cues on the walls ensured consistent spatial relationships that defined the boundaries of the recording room across days.

### Surgery

Rats were surgically implanted with tetrode arrays (twisted sets of four 12 micrometer tungsten wires or 17 micrometer platinum-iridium wires) fitted to custom-fabricated microdrives that allowed movement in 40μm increments. Each microdrive contained 4 tetrodes. Rats were implanted with 3 microdrives (2-3 bilateral RSC, 1 HPC, depending on animal). Rats were anesthetized with isoflurane and positioned in a stereotaxic device (Kopf Instruments). Following craniotomy and resection of dura mater overlying the retrosplenial cortex, microdrives were implanted relative to bregma (A/P −5.8 mm, M/L ± 0.7-1.2 mm, D/V −0.5 mm, 10-12° medial/lateral angle). 3 animals received a unilateral HPC microdrive targeted to the CA1 sub-region (target coordinates relative to bregma, A/P −3.8mm and M/L ± 2.3mm, D/V −0.5mm).

### Recordings

Each microdrive had one or two electrical interface boards (EIB-16, Neuralynx) individually connected to amplifying headstages (20X, Triangle Biosystems). Signals were initially amplified and filtered (50×, 150Hz high-pass) on the way to an acquisition computer running Plexon SortClient software. Here the signal was digitized at 40kHz, filtered at 0.45-9 kHz and amplified 1-15X (to reach a total of 1,000-15,000X). Electrodes were moved ventrally (40μm) between recordings to maximize the amount of distinct units collected for each animal.

Animal position was tracked using a camera set 10ft above the recording room floor. Plexon’s CinePlex Studio software was utilized to separately detect blue and red LED lights. Lights sat approximately 4.5 cm apart and were positioned perpendicular to the length of the animal’s head. Recordings lasted approximately 20-45 minutes, the amount of time needed for the animal to complete an absolute minimum of 5 ballistic runs for plus and/or ring track conditions.

### Unit isolation and sort quality

Single-units were identified using Plexon OfflineSorter software. Primary waveform parameters utilized were peak height, peak-valley, energy, and principal components. To assess sort quality, custom MATLAB software was developed to compute isolation distance and L-Ratio metrics for each unit^53,54^.No units were excluded on the basis of their cluster quality scores. Instead, we show the entire distribution of L-ratio and Isolation Distance values to demonstrate that neurons exhibiting spatial periodicity and distance encoding had a range of cluster quality scores, from average to extremely well isolated (**Figure S1**).

The only RSC units that were excluded from analyses on the tracks were those that did not exhibit peak activation of at least 3Hz for a single track bin and those that were statistically identified as head direction neurons. HPC neurons were excluded if they had mean activity across all track sessions greater than 8Hz (i.e. fast-firing, putative interneurons), no bins in which the firing rate dropped to 0Hz (i.e. fast-firing, putative interneurons), and no bins exhibiting a peak firing rate amplitude above 3Hz.

### Histology

Animals were perfused with 4% paraformaldehyde under deep anesthesia. Brains were removed and sliced into 50μm sections and Nissl-stained to identify the trajectory and depth of electrode wires in RSC and HPC. RSC was defined in accordance with our previous work^5^ in the region as well as the Paxinos and Watson^55^ and Zilles^56^ atlases. The boundary of RSC was considered to be approximately 1-2mm lateral of the midline depending on both the rostrocaudal and dorsoventral position of the tetrode. Ventrally, the lateral edge was defined by the transition from retrosplenial cortex to the subiculum. All tetrodes were determined to be within the bounds of RSC. Documented microdrive depth across recordings and final electrode depth observed in histology were compared and found to be compatible in all cases.

### Identification of ballistic track traversals

Position tracking data was pulled into a custom MATLAB guided user interface. From trial-to-trial, runs in which the animal moved uninterruptedly across the track were identified as ‘clean’ traversals and pulled out for subsequent analysis. A minimum of 5 uninterrupted traversals was required for a recording session to be included in subsequent analyses. The average number of traversals on the plus and ring tracks were 14 and 13, respectively.

### Track linearization and firing rate calculation

The space of the track was linearized to analyze spatial dependency of neural activity. Custom MATLAB software was utilized to generate a spatial template match for the average movement of the animal through pixel space along the track. All full, half, and ring trials were plotted independently and the coordinates of track start, end, and apex of all turns for each route were identified (in pixel space) based on each animal’s unique and stereotyped movement through the track. Between behavioral epochs (start, end, and turn apices) a series of template bins was generated with a spacing of 10 pixels (approx. 3.5 cm). Each session was linearized by fitting both position and neural data acquired on each trial to template space. Firing rates were then computed for each template bin by dividing the total number of spikes by the time of occupation. Activity patterns were smoothed with a narrow Gaussian filter (7 cm s.d.).

Like the linear firing ratemaps, two dimensional firing ratemaps were constructed for each track running session individually and during ballistic track traversals only. Tracking data was binned into 3.5cm × 3.5cm spatial bins and the animal’s occupancy in seconds for each was determined as well as the corresponding number of spikes. Firing rates were computed by dividing the number of spikes in each bin by the total time in seconds that each bin was occupied. Raw two dimensional ratemaps were smoothed with a Gaussian kernel (7 cm s.d.).

### Directional tuning sensitivity

Head direction (HD) was calculated as a function of the angle of two tracking LEDs placed laterally on the rat’s head. For each neuron, the firing rate for each heading direction, discretized to 5° bins, was calculated as the total number of spikes divided by the total time in seconds that the heading bin was occupied. Two head direction tuning curves were computed for each neuron, for the first and second half of the recording separately. Neurons with strong directional tuning that was reliable across these two head-direction tuning vectors were eligible to be classified as HD neurons. The length and mean direction of the resultant vector, as well as a Rayleigh test for non-uniformity were calculated for both tuning curves using the Circular Statistics toolbox for MATLAB^57^. RSC cells that were statistically non-uniform in their firing rate as a function of heading direction, exhibited consistent mean directional tuning, and had large resultant lengths (>0.2) across both halves of the recording were determined to be head-direction neurons (see Fig. S1). Head direction neurons were removed from all analyses on the track.

### Correlative route position and distance reconstructions

To determine the extent to which route positions were encoded across the entire RSC population, a correlative route position reconstruction was conducted for plus track traversals. Individual mean firing rate profiles were computed using data collected on odd and even trials separately. Correlation matrices were constructed with these spatial activation profiles to estimate the animal’s position on even trials from data collected on odd trials. In the correlation matrices of **Figure 1C** and **Figure 1D**, each row corresponds to the correlation of the odd-trial population vector at a single track location (i.e. a template bin) with the even-trial population vector at every track location (i.e. all template bins). For each row of the matrix, the column with the maximal correlation is the predicted position of the rat on even trials, because it is the position that yielded the greatest similarity to the odd-trial population rate vector for the track bin corresponding to that row of the matrix.

A similar correlative reconstruction process was computed for the distance reconstructions shown in **Figure 6D**. Here, instead of correlating population rate vectors taken from odd and even trials we correlate population rate vectors taken from the first half of the symmetrical rate vectors (**see Figure 1C, 1F**) against those taken from the second half. In this manner, the estimate is not of the spatial location of the rat, but rather the animal’s distance from the point of symmetry. Again, the estimate is quantitatively the maximum correlation in each row.

### Spatial and Behavioral Generalized Linear Models

A series of spatial GLMs were implemented to assess the presence of spatial periodicity in the activation patterns of RSC neurons on the plus and ring tracks. To begin, a total of 6 spatial predictors were constructed using paired sine and cosine functions. Spatial predictors were generated to split the full space of the plus into multiples of 12, as there were 12 total segments on the plus. For a given predictor, the total number of complete cycles of the sine and cosine functions would take on discrete values: twelve, six, four, three, two, or one. The total number of complete cycles within the predictor would correspond to fragmentation of the space into single segments (twelfths), sixths, quarters, thirds, halves, and full (**Figure 2A**). All fragmentations were anchored relative to the first bin in linear space. Using this method, corresponding linear track positions within each fragment (i.e. the first bin of the first half of the track and the first bin of the second half of the track) have identical predictor values defining spatial position within the fragmentation. The same spatial predictors that fragmented the plus track into multiples of 12 were implemented in creating model fits to firing rate profiles taken from ring traversals.

Following construction of spatial predictors, a complete generalized model (cGLM) was constructed for the max-normalized mean firing rate profile in linearized track space for each neuron using all spatial predictors (‘glmfit’ with identity link function in MATLAB). Regression coefficients were evaluated (‘glmval’ in MATLAB) to construct the predicted firing rate profile for the neuron and the fit between the actual and predicted firing rate profiles was assessed via normalized mean squared error (NMSE, ‘goodnessOfFit’ function in MATLAB). To test the impact of each spatial predictor, partial GLMs (pGLM) were fit dropping each predictor individually and computing the model fit. Kruskal-Wallis tests with post hoc Bonferonni corrections were conducted on distributions of fitness values for the pGLMs relative to the cGLM.

To test the significance of individual predictors for individual neurons, cGLMs and pGLMs were constructed for trial firing rate vectors individually. Thus, for a single neuron, a distribution of trial fitness values was generated for cGLMs and all pGLMs which could then be tested for significance. Neurons were considered conjunctive in their periodic structure if they had significant decrements to model fit for multiple pGLMs.

cGLMs and pGLMs were also generated using behavioral (egocentric) predictors in place of the spatial predictors described above. Behavioral predictors utilized were linear speed, linear acceleration, and angular velocity profiles. Again, these models were assessed using the mean firing rate profiles of individual neurons and mean behavioral predictors, or individually for each trial. This method was applied to both plus and ring track traversals.

### Identification of symmetry points in spatial firing rate profiles

Points of symmetry (SP) within individual spatial firing rate profiles were identified for analysis of distance encoding in RSC ensembles. Identifying SPs was an iterative process wherein the spatial firing rate profile for each neuron was circularly rotated (‘circshift’ in MATLAB) a single position and the rotated vector was smoothed with a box car filter spanning 5 template bins (17.5 cm). The vectors were smoothed following rotations in an effort to reduce any drastic jumps in firing rate that may occur between the edges (first and final template bins) of the firing rate profile. After smoothing, the rotated vector was reflected from the center point and the NMSE was computed between the two halves and stored. This process continued until the vector had been rotated a full 360°. The rotation that produced the maximal fit between the reflected halves was determined to be the SP. The rotated vector that corresponded to the SP was stored for further analysis.

### Distance reconstruction with random sub-sampling of population vectors from RSC and HPC

Distance estimates were computed by sub-sampling N-matched, randomly-selected sub-populations of RSC and HPC neurons. Population vectors were extracted from rotated, symmetrical, firing rate profiles from both RSC and HPC. Population vectors were sampled in multiples of 5, beginning at 5 neurons and ending at 140. For each vector size, neurons were resampled 100 times and a distance estimate was computed.

---

**Figure S1.**
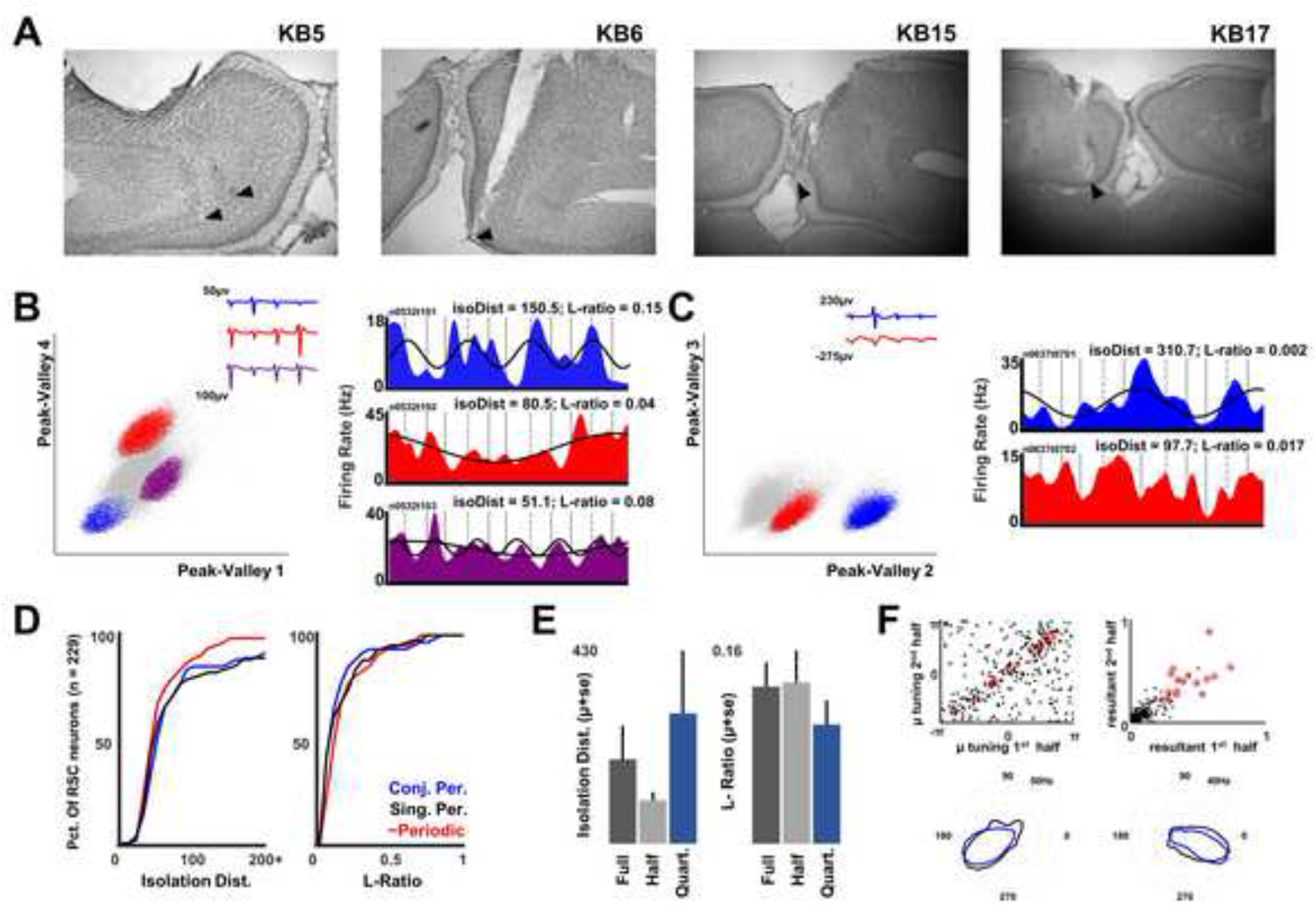
Representative histology, sort quality, and head-direction assessment. **A.** Final placement of electrode tracts in RSC for example rats. **B and C.** Left, example unit isolation and waveforms for a single tetrode. Right, corresponding spatial firing rate profiles for each neuron. Above each graph, scores for the cluster quality metrics isolation distance and L-ratio. **D.** Cumulative distribution of sort quality metrics. Left, isolation distance. Right, L-ratio. For each plot, the cumulative percentage of neurons with sort quality values less than each value on the x-axis is shown in colored lines. In red, cumulative density of neurons that did not exhibit sensitivity to spatially periodic predictors. In black, cumulative density of neurons that exhibited sensitivity to a single spatial predictor. In blue, cumulative density of neurons that exhibited sensitivity to multiple spatial predictors. **E.** Mean (± s.e.) isolation distance and L-ratio for neurons that were sensitive to full, half, and quarter spatial periodicities. **F.** Method for assessment of significant directional tuning. For each neuron, the mean directional tuning vector and resultant for the first and second half of the recording session were computed individually. Between the two blocks, RSC neurons that had mean tuning differences less than 0.35 radians, resultants greater than 0.2, and exhibited significantly non-uniform firing rate distributions as a function of heading (Rayleigh test, p<0.05) were identified as head-direction neurons and removed from further analyses. Two example neurons are shown in the bottom row of the graph with the tuning curves from the two halves of the recording session shown in blue and black.

**Figure S2.**
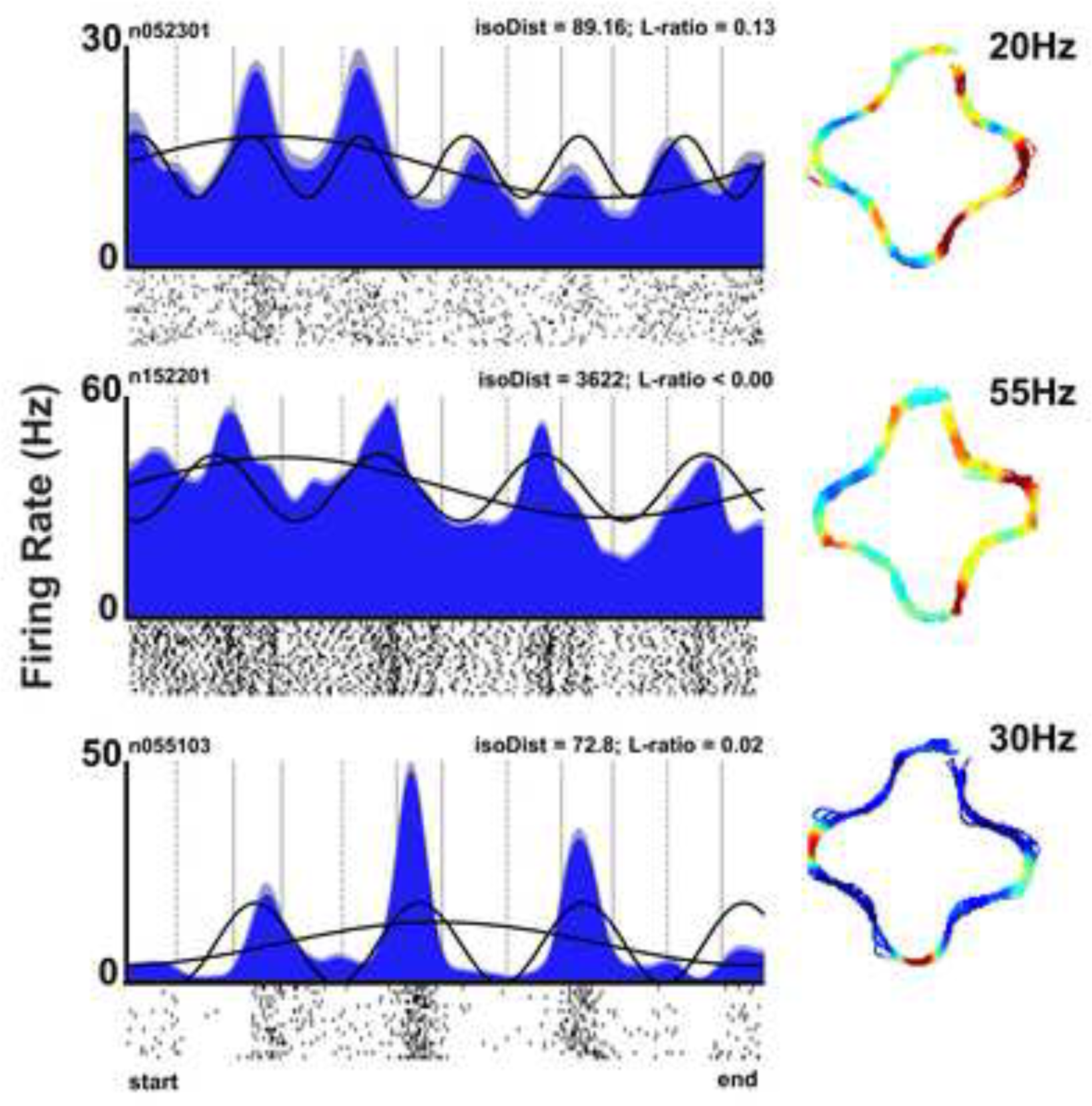
RSC neurons exhibit conjunctive sensitivity to multiple sub-route and full route spatial oscillations. Mean spatial firing rate profiles (± s.e.) for three example neurons. Overlayed in black, iGLM fits for individual spatial predictors that significantly modulated each neuron (as assessed with tests of trial-by-trial pGLMs versus cGLMs). Above each plot, sort quality metrics for each neuron demonstrate that conjunctive sub-route sensitivity was not the product of poor unit isolation. Below each plot, corresponding spike trains across trials. Right, corresponding two dimensional ratemap with max firing rate indicated (blue indicates zero firing).

**Figure S3.**
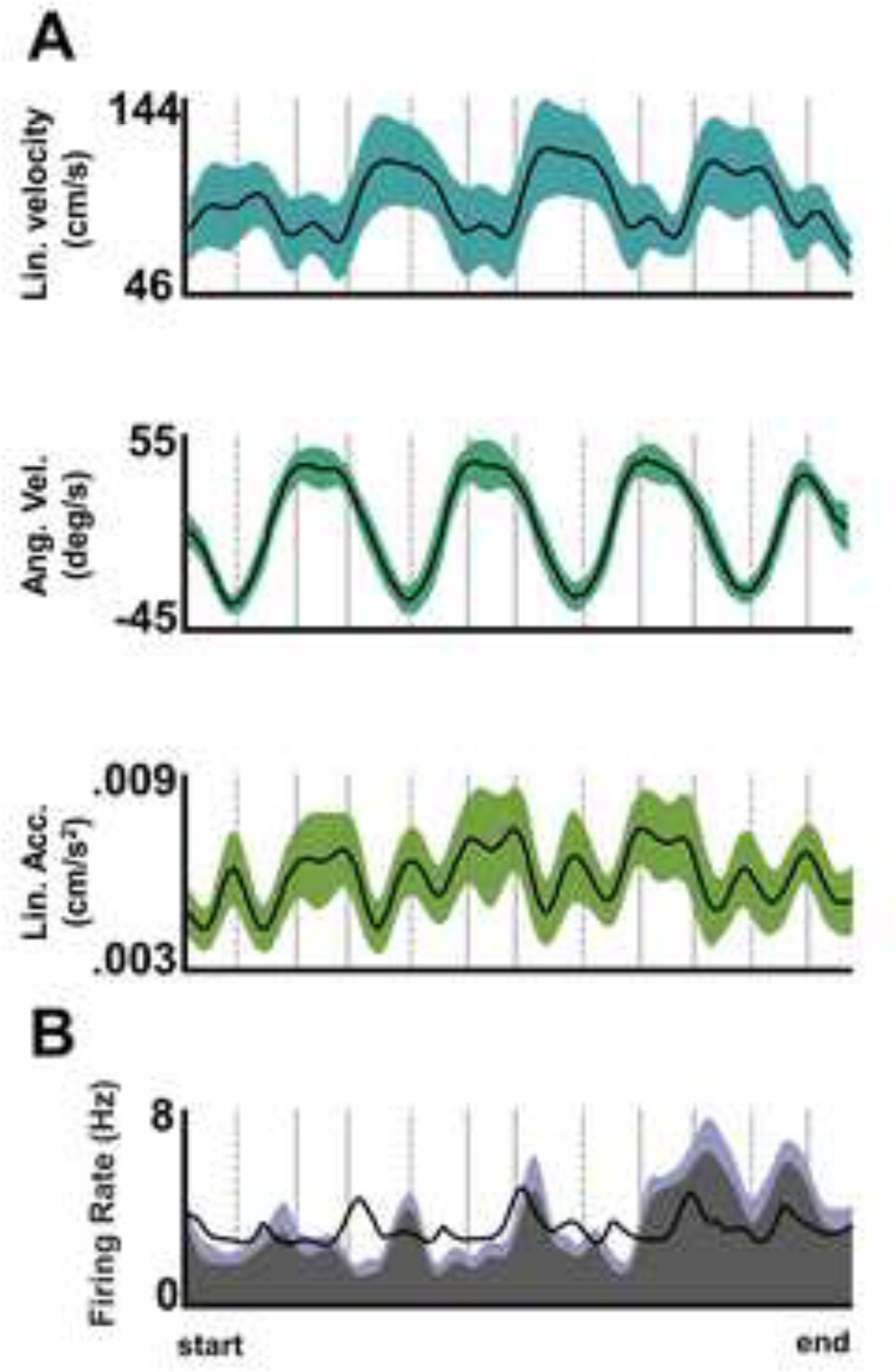
Description of movement variables utilized in egocentric GLM during plus track running. **A.** Mean (± s.d.) linear movement variables across all recording sessions (n = 91) and animals for the plus track. Top, linear velocity; Middle, angular velocity; Bottom, linear acceleration. **B.** Mean spatial firing rate profile (± s.d.) for a neuron during a single recording session. In black, complete model fit to the activation profile of the neuron using all three movement variables from the corresponding recording of the neuron.

**Figure S4.**
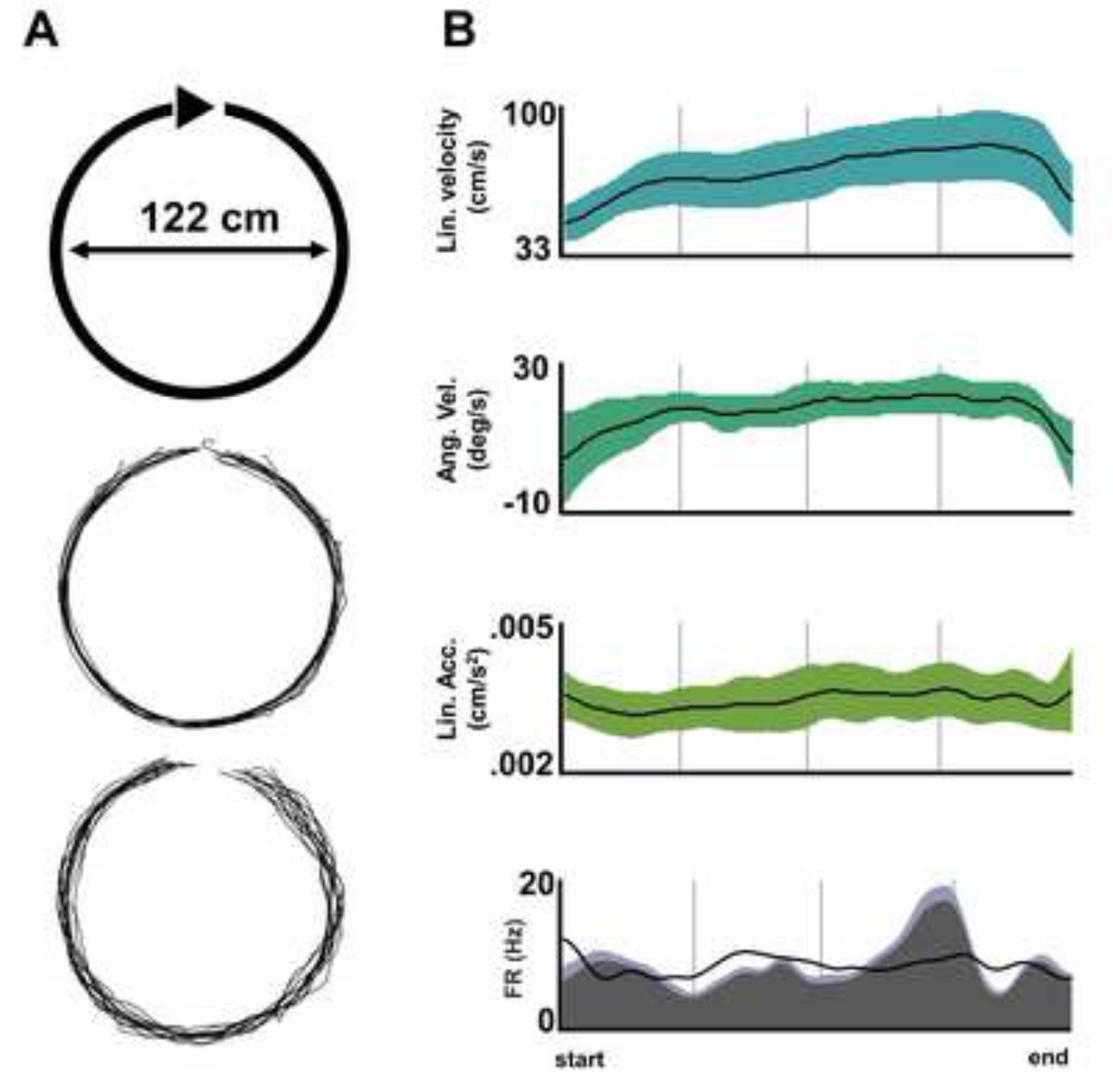
Description of movement variables utilized in egocentric GLM during ring track running. **A.** Top, schematic of ring track. Bottom two plots, example positional tracking from two rats and two ring track sessions. **B.** Mean (± s.d.) linear movement variables across all recording sessions (n = 35) and animals for the ring track. Top, linear velocity; Middle, angular velocity; Bottom, linear acceleration. Bottom plot, mean spatial firing rate profile (± s.d.) for a neuron during a single recording session. In black, complete model fit to the activation profile of the neuron using all three movement variables from the corresponding recording of the neuron.

